# Cis-Compound Mutations are Prevalent in Triple Negative Breast Cancer and Can Drive Tumor Progression

**DOI:** 10.1101/085316

**Authors:** Nao Hiranuma, Jie Liu, Chaozhong Song, Jacob Goldsmith, Michael O. Dorschner, Colin C. Pritchard, Kimberly A. Burton, Elisabeth M. Mahen, Sibel Blau, Francis M. Senecal, Wayne L. Monsky, Stephanie Parker, Stephen C. Schmechel, Stephen K. Allison, Vijayakrishna K. Gadi, Sophie R. Salama, Amie J. Radenbaugh, Mary Goldman, Jill M. Johnsen, Shelly Heimfeld, Vitalina Komashko, Marissa LaMadrid-Hermannsfeldt, Zhijun Duan, Steven C. Benz, Patrick Soon-Shiong, David Haussler, Jingchun Zhu, Walter L. Ruzzo, William S. Noble, C. Anthony Blau

## Abstract

About 16% of breast cancers fall into a clinically aggressive category designated triple negative (TNBC) due to a lack of ERBB2, estrogen receptor and progesterone receptor expression^1-3^. The mutational spectrum of TNBC has been characterized as part of The Cancer Genome Atlas (TCGA)^4^; however, snapshots of primary tumors cannot reveal the mechanisms by which TNBCs progress and spread. To address this limitation we initiated the Intensive Trial of OMics in Cancer (ITOMIC)-001, in which patients with metastatic TNBC undergo multiple biopsies over space and time^5^. Whole exome sequencing (WES) of 67 samples from 11 patients identified 426 genes containing multiple distinct single nucleotide variants (SNVs) within the same sample, instances we term Multiple SNVs affecting the Same Gene and Sample (MSSGS). We find that >90% of MSSGS result from cis-compound mutations (in which both SNVs affect the same allele), that MSSGS comprised of SNVs affecting adjacent nucleotides arise from single mutational events, and that most other MSSGS result from the sequential acquisition of SNVs. Some MSSGS drive cancer progression, as exemplified by a TNBC driven by FGFR2(S252W;Y375C). MSSGS are more prevalent in TNBC than other breast cancer subtypes and occur at higher-than-expected frequencies across TNBC samples within TCGA. MSSGS may denote genes that play as yet unrecognized roles in cancer progression.

---

The Intensive Trial of OMics in Cancer (ITOMIC)-001 (ClinicalTrials.gov Identifier: NCT01957514) enrolls patients with metastatic TNBC who have not received platinum based therapies and are scheduled to receive cisplatin^5^. Up to seven different metastatic sites are biopsied prior to cisplatin, following discontinuation of cisplatin, and following subsequent therapies. Additional samples are accessed as archival tissues, as leftovers following clinically indicated procedures, and from tissues taken at autopsy. Samples are selected for sequencing based on specimen size and tumor content (Fig. 1a). Results from 11 of the first 12 subjects are presented here because low tumor purities in Subject 08 precluded analysis. We performed WES of germline DNA and 67 tumor samples (**Extended Data Table 1**), including transiently cultured cells derived from a malignant pleural effusion in Subject 01 and two highly enriched samples of circulating tumor cells (CTCs) obtained following leukapheresis in Subject 02^5^. Whole genome sequencing (WGS) was also performed in 39 samples. The number of assessed samples per patient ranged from 2 to 16 (median 7), and the number of assessed time points ranged from 1 to 6 (median 3) (Table 1).

**Figure 1.**
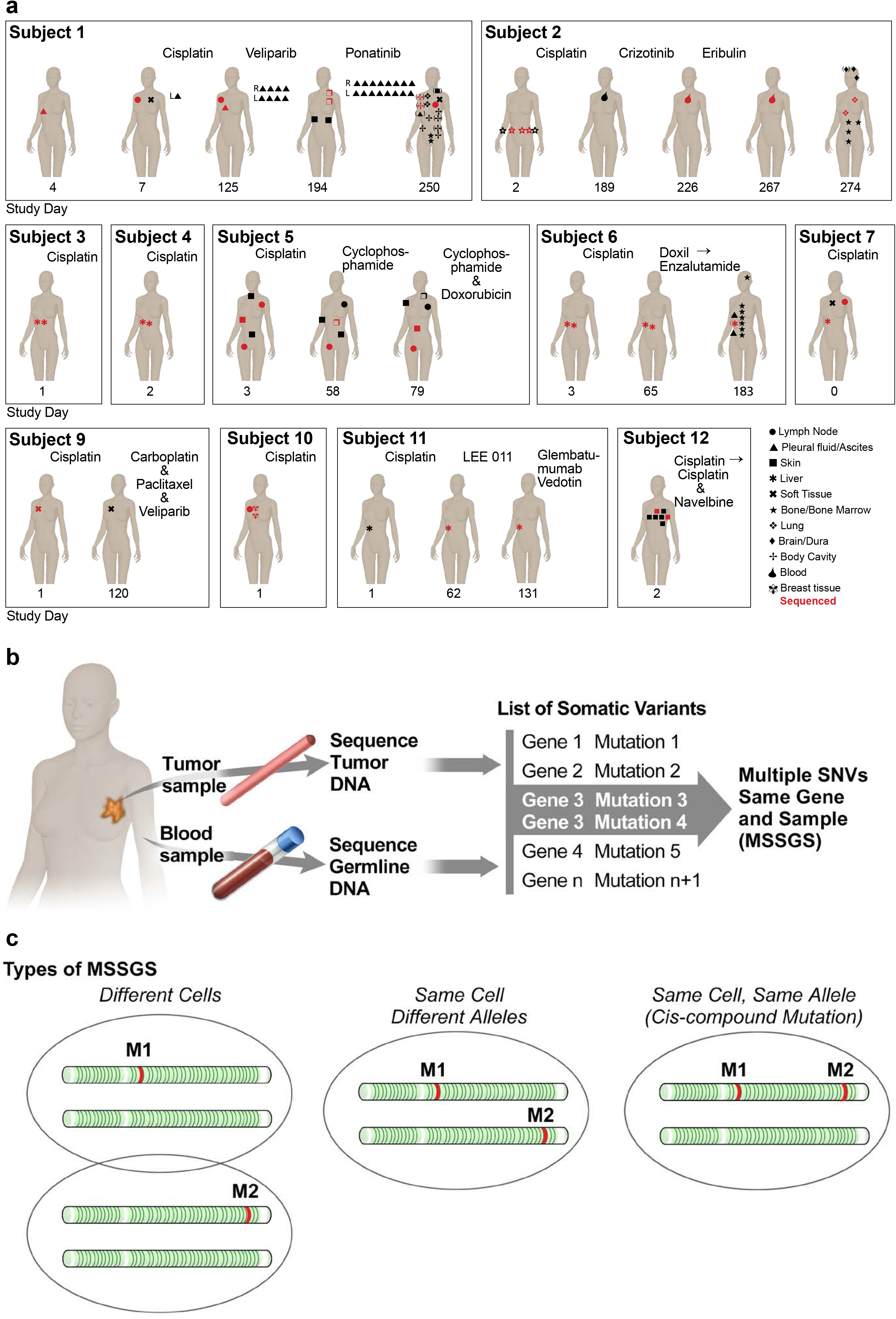
Schematic depiction of tumor samples collected and analyzed as part of ITOMIC-001 and description of MSSGS. (**a**) Schematic depiction of biopsies performed (black symbols) and sequenced (red symbols) in 11 of the first 12 subjects enrolled in ITOMIC-001. All treatments during participation in the study are as described. Subject 8 is not depicted because low tumor purities precluded analysis. (**b**) MSSGS are defined by the presence of two or more somatic SNVs occurring within the same gene and tumor sample. (**c**) MSSGS may arise from: SNVs occurring in different populations of tumor cells (left), SNVs affecting different alleles within the same cells (trans-compound mutation) (middle), SNVs affecting the same allele (cis-compound mutation) (right).

**Table 1.**
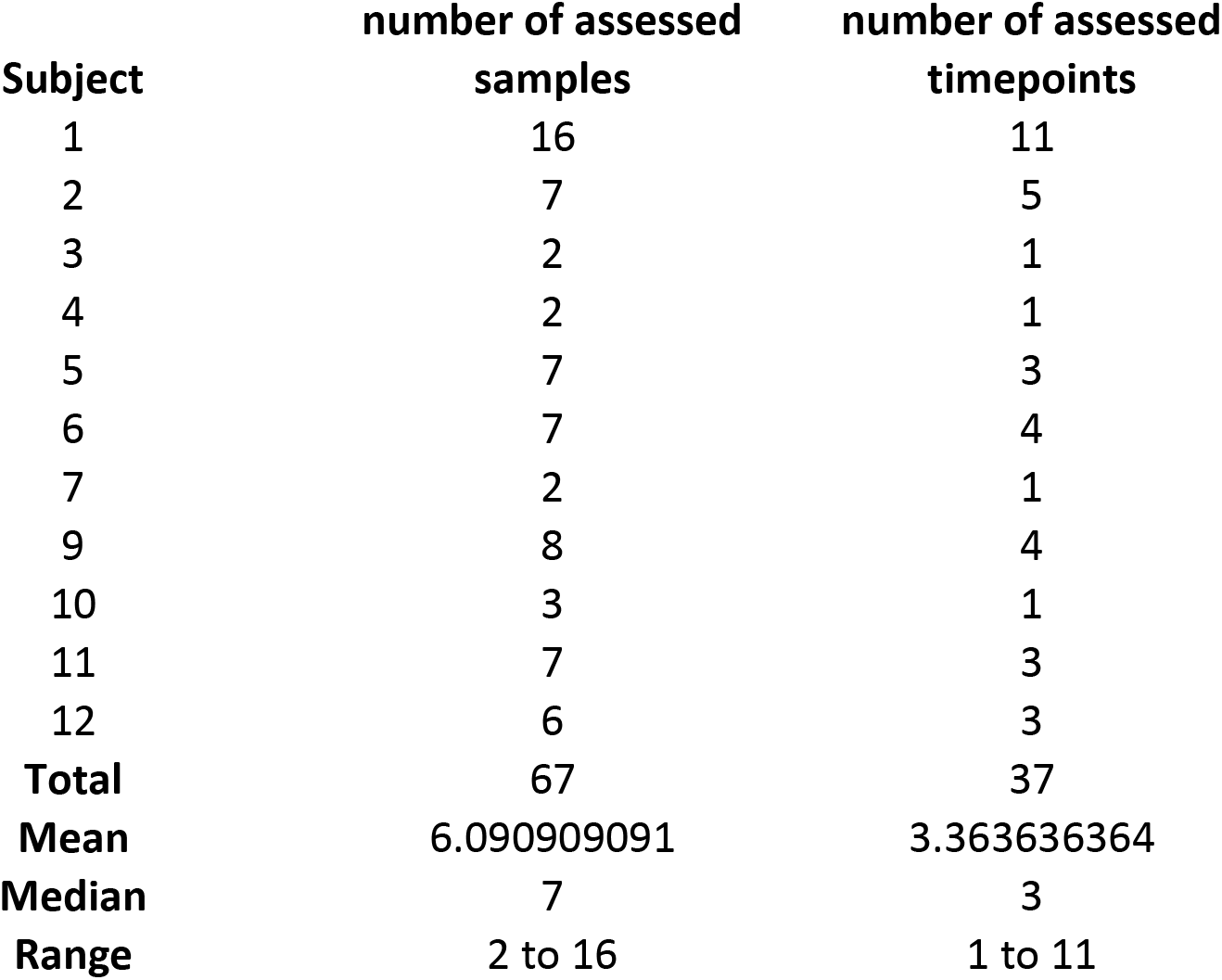
Numbers of assessed patients, samples and timepoints.

| **Subject** | **number of assessed**<br>**samples** | **number of assessed**<br>**timepoints** |
| --- | --- | --- |
| 1 | 16 | 11 |
| 2 | 7 | 5 |
| 3 | 2 | 1 |
| 4 | 2 | 1 |
| 5 | 7 | 3 |
| 6 | 7 | 4 |
| 7 | 2 | 1 |
| 9 | 8 | 4 |
| 10 | 3 | 1 |
| 11 | 7 | 3 |
| 12 | 6 | 3 |
| **Total** | 67 | 37 |
| **Mean** | 6.090909091 | 3.363636364 |
| **Median** | 7 | 3 |
| **Range** | 2 to 16 | 1 to 11 |

Our analysis focused on somatic single nucleotide variants (SNVs), which comprise the majority of mutations in breast cancer^4^,^6^,^7^. WES identified a total of 8449 SNVs, of which 7067 occurred within 5136 protein coding genes, including 43 breast cancer driver genes (0.84%)^7^ (data not shown). Across all 67 samples, 1403 genes were affected by >1 SNV. Remarkably, 426 genes were found to contain >1 SNV within the same tumor sample (**Extended Data Table 1**), and we designated these instances Multiple SNVs affecting the Same Gene and Sample (MSSGS) (Fig. 1 b).

MSSGS were observed in 65 of 67 samples, with a median of 8 and a range of 0 to 133 MSSGS per sample (**Extended Data Table 1**). The distribution of median transcript sizes was no larger for MSSGS than for genes affected by single SNVs^8^ (Extended Data Fig. 1). Concomitant WGS in 39 of 65 samples independently confirmed 48 of 150 evaluable MSSGS (32%), with most of the unconfirmed cases being attributable to the relatively lower read counts associated with WGS (**Extended Data Table 2 and Supplementary Methods**), although a fraction of MSSGS (11% - 15%) may represent artifacts associated with WES (**Supplementary Methods**). While somatic variants are unevenly distributed across cancer genomes^8^, we found similar mutation densities in regions surrounding MSSGS compared to isolated SNVs (Fig. 2a) with the exception of an MSSGS involving *MUC4*, which fell within a region of kataegis^9^ (Fig. 2b).

**Figure 2.**
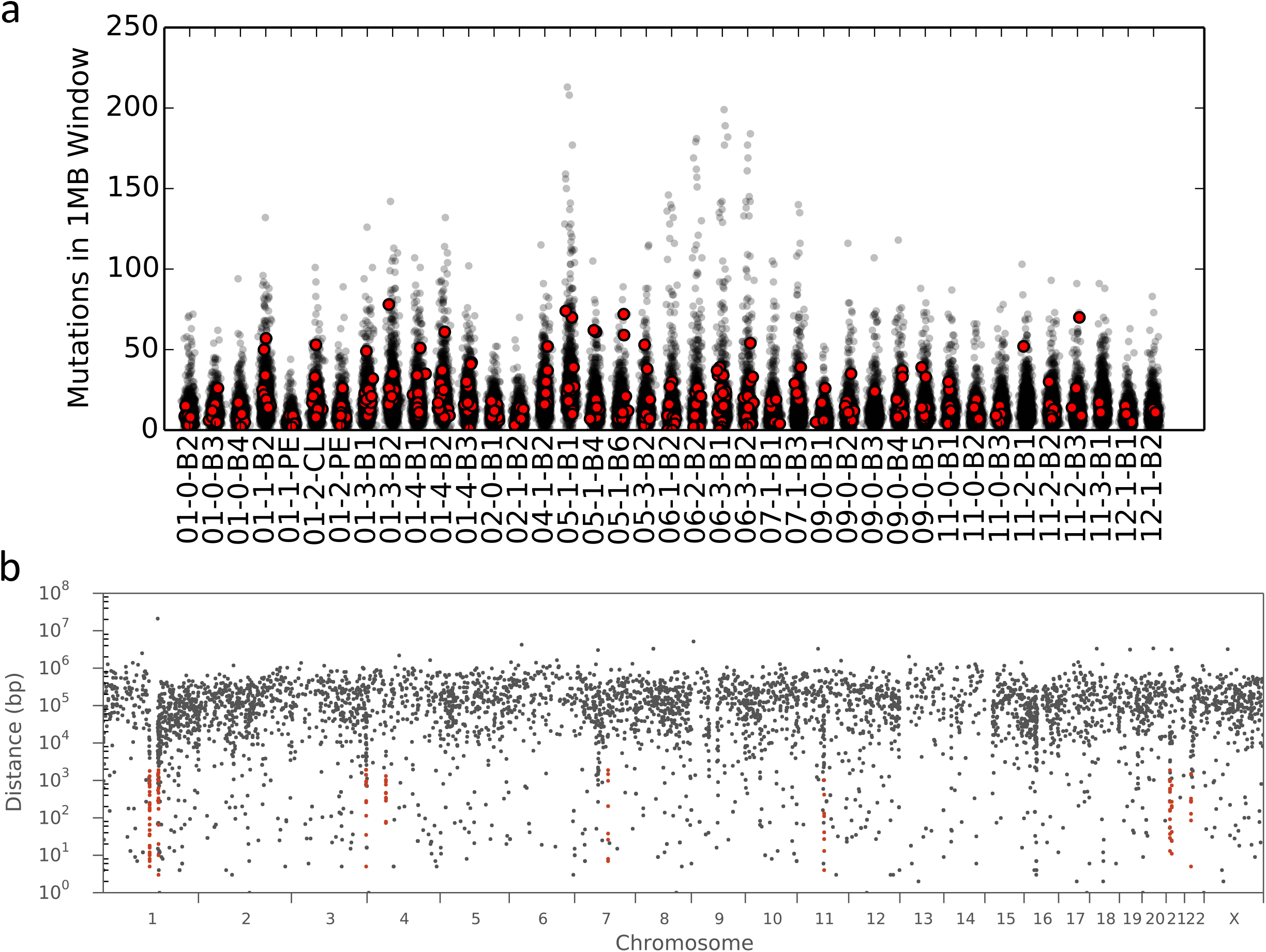
MSSGS are not associated with an increase in regional mutation
density. (**a**) Y axis indicates numbers of SNVs in 1MB windows across the genome (gray circles) and surrounding MSSGs (red circles) for each sample examined by WGS (X axis). (**b**) Mutational density plot for sample 11-2-B3. The Y axis indicates the distance (in log scale) between two neighboring SNVs, and the X axis indicates the chromosomal position of each SNV-pair. SNVs in regions of kataegis are colored with red circles and the position of *MUC4* is labeled in blue.

MSSGS can arise when two or more SNVs affect: i) the same gene in different tumor cells, ii) different alleles in the same cell, or iii) the same allele in the same cell (cis-compound mutation) (Fig. 1c). To assess the frequency with which SNVs contributing to MSSGS co-localize to form a cis-compound mutation we cloned and sequenced tumor DNA from eight patients, revealing a very high frequency of cis-compound mutations (12 of 13 evaluable MSSGS - 92%) (Table 2). Haplotype phasing^10^ of 407 MSSGS similarly indicated that >90% were attributable to cis-compound mutations (**Extended Data Table 3**). This tendency for SNVs contributing to MSSGS to affect the same allele may result from a DNA strand bias in susceptibility to point mutations^11^, or from a selection of genes bearing cis-compound mutations.

**Table 2.** Results from cloning and sequencing 15 MSSGS-containing fragments. 12 of 13 evaluable MSSGS were associated with cis-compound mutations. Clonal analysis of 15 MSSGS. Estimated tumor content is based on tumor content in formalin fixed half of each biopsy specimen. CTCs: Circulating Tumor Cells; REF: Number of sequenced DNA fragments containing the reference (non-mutated) sequence; nt: Nucleotide; chr: Chromosome; aa: Amino Acid; * *ZZZ3* contained 3 SNVs, *F8* contained four MSSGS but only two were interrogated. All other MSSGS shown here contained two SNVs; ** APOBR could not be evaluated due to sequencing artifacts; Estimate following column enrichment and FACs sorting of CTCs; § Touch prep of the snap frozen tissue used for sequencing demonstrated numerous tumor cells.

| # | Subject | Estimated Tumor Content (Disease Site) | Gene | WES |  | DNA fragment sequencing |  |  |  |  |
| --- | --- | --- | --- | --- | --- | --- | --- | --- | --- | --- |
|  |  |  |  | SNV1<br>nt change<br>chr position<br>aa change<br>VAF | SNV2<br>nt change<br>chr position<br>aa change<br>VAF | REF<br>% | SNV1<br>% | SNV2<br>% | SNV1+<br>SNV2<br>% | Clones<br>Tested<br>(Assessable) |
| 1 | 06 | 85%<br>(Liver) | *ABHD16Ac* | C-**T**<br>6:31659632<br>Q230H<br>0.473 | C-**G**<br>6:31659583<br>E247K<br>0.424 | 64% | 4% | 2.7% | 28% | 75<br>(94) |
| 2 | 01 | 75%<br>(Pleura) | *ADAM33* | C-**A**<br>20:3652402<br>Q577H<br>0.131 | T-**A**<br>20:3652403<br>Q577L<br>0.130 | 96.8% | 0% | 0% | 3.2% | 94<br>(94) |
| 3 | 06 | 70%<br>(Liver) | *APOBR* ** | G-**C**<br>16:28507445<br>E361D<br>0.161 | G-**T**<br>16:28507452<br>G364W 0.107 | 100% | - | - | - | 4<br>(94) |
| 4 | 10 | 60%<br>(Breast) | *C20orf26* | G-**C**<br>20:20037448<br>NA<br>0.152 | G-**C**<br>20:20037264<br>NA<br>0.11 | 81% | 6.6% | 7.7% | 4.4% | 91<br>(94) |
| 5 | 01 | 0% §<br>(Skin) | *CBR4* | C-**A**<br>4:169923348<br>V137F<br>0.228 | T-**A**<br>4:169923351<br>I136F<br>0.145 | 100% | 0% | 0% | 0% | 92<br>(92) |
| 6 | 04 | 70%<br>(Liver) | *CCDC171* | T-**G**<br>9:15744665<br>K815T<br>0.11 | A-**C**<br>9:15744263<br>NA<br>0.106 | 94.5% | 2.2% | 1.1% | 2.2% | 92<br>(94) |
| 7 | 01 | 0% §<br>(Skin) | *F8* * | A-**T**<br>X:154157659<br>L1469*<br>0.107 | T-**A**<br>X:154157737<br>K1443I<br>0.101 | 96% | 1% | 1% | 3% | 93<br>(94) |
| 8 | 04 | 65%<br>(Liver) | *FLG2* | A-**T**<br>1:152329531<br>L244*<br>0.259 | C-**G**<br>1:152329951<br>R104P<br>0.209 | 72% | 14% | 3% | 1% | 93<br>(94) |
| 9 | 06 | 70%<br>(Liver) | *MGA* | G-**C**<br>15:42042226<br>E2141Q<br>0.374 | G-**C**<br>15:42041736<br>Q1977H<br>0.385 | 50 % | 23% | 17% | 10% | 87<br>(94) |
| 10 | 05 | 30%<br>(Skin) | *TCTE1* | C-**G**<br>6:44254146<br>C134S<br>0.122 | A-**G**<br>6:44254159<br>Y130H<br>0.128 | 95% | 2% | 3% | 0% | 93<br>(94) |
| 11 | 01 | 75%<br>(Pleura) | *TMPRSS13* | T-**C**<br>11:117789342<br>Q78R<br>0.165 | G-**C**<br>11:117789345<br>A77G<br>0.120 | 48% | 2% | 0% | 49% | 91<br>(94) |
| 12 | 06 | 95%<br>(Liver) | *DMPK* | C-**T**<br>19:46283223<br>R32K<br>0.32 | C-**T**<br>19:46283028<br>NA<br>0.331 | 70% | 7% | 9% | 14% | 100<br>(101) |
| 13 | 02 | 100%¶<br>(CTCs) | *ROBO4* | G-**C**<br>11:124754988<br>Q984E<br>0.606 | G-**C**<br>11:124755134<br>S935*<br>0.555 | 28% | 9% | 3% | 60% | 102<br>(103) |
| 14 | 02 | 100%¶<br>(CTCs) | *ZZZ3* * | TCC-**AAT**<br>1:78045223<br>1:78045224<br>1:78045225<br>R690N, T691S<br>0.41, 0.4, 0.4 |  | 57% | 0% | 0% | 43% | 100<br>(101) |
| 15 | 07 | 90%<br>(Liver) | *FANCC* | G-**C**<br>9:98011451<br>F41L<br>0.295 | G-**C**<br>9:98011537<br>Q13E<br>0.22 | 62% | 13% | 6% | 19% | 102<br>(103) |

CTCs: Circulating Tumor Cells
nt: Nucleotide
chr: Chromosome
aa: Amino Acid

Assessing SNVs across different tumor samples from the same patient provides insight into the spatial and temporal origins of MSSGS. 130 of 426 MSSGS (~30.5%) were detected in at least two tumor samples from the same patient. Among these repeatedly detected MSSGS, three contiguous SNVs affecting ZZZ3 were all detected in the same 4 of 7 samples from Subject 02 at nearly identical frequencies, suggesting their presence in the same cells (**Extended Data Table 1**), and a similar pattern was observed for *ADAM33*, which was detected in the same 3 of 16 samples from Subject 01 (Fig. 3a, **Extended Data Table 1**). These patterns align with results from cloning and sequencing these MSSGS fragments (Table 2), which showed that either all SNVs contributing to the cis-compound mutation were observed or none were observed, consistent with their emergence from single mutational events.

**Figure 3.**
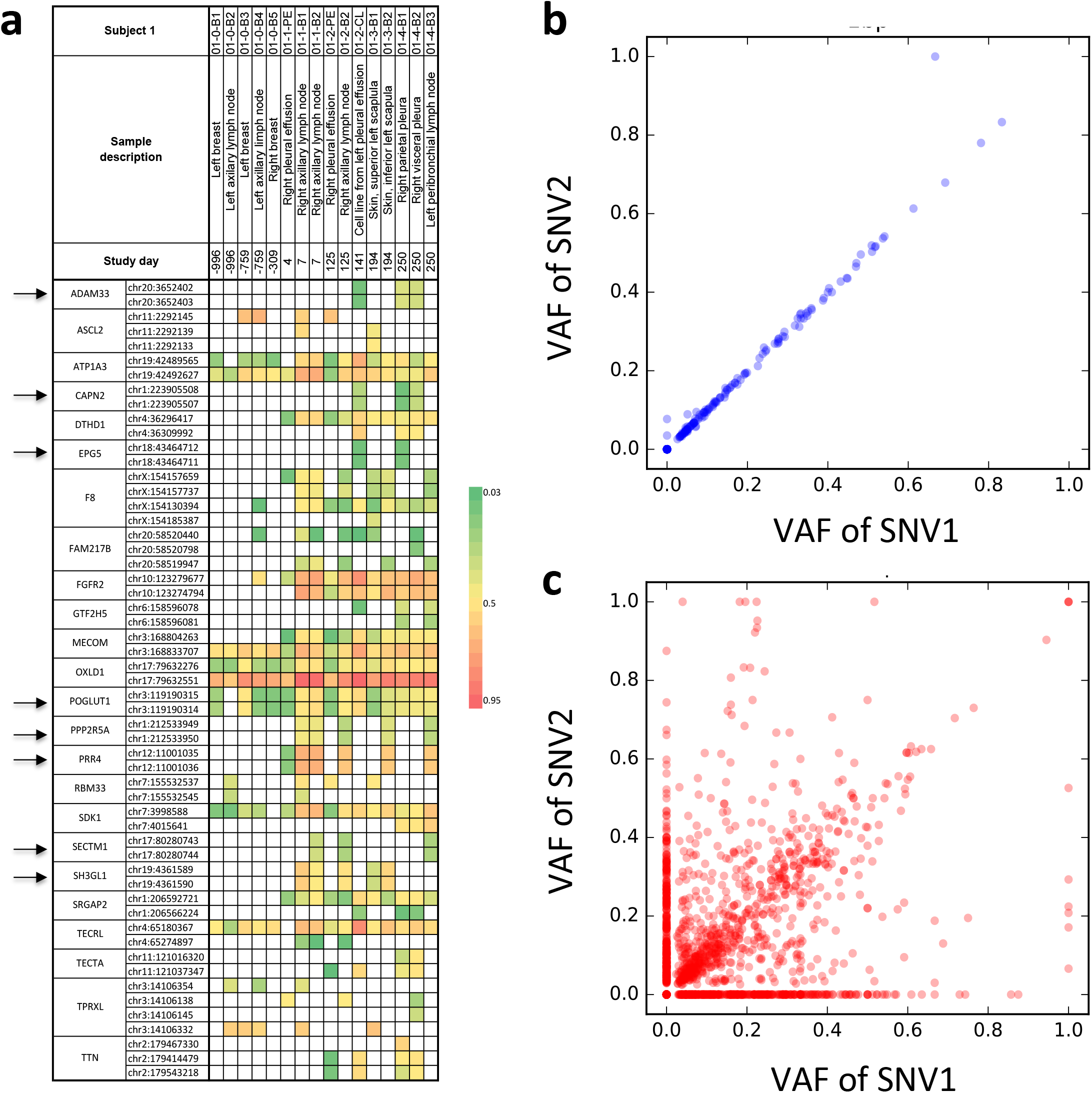
MSSGS comprised of SNVs that affect consecutive nucleotides appear abruptly, whereas most other MSSGS develop over time. (**a**) Appearance of SNVs contributing to MSSGs in samples collected over time from Subject 1. Genomic positions of SNVs are indicated. Colors depict VAFs as indicated in the key. Note that MSSGS comprised of SNVs affecting adjacent nucleotides (arrows) consistently appear together and at very similar VAFs. (**b**) VAFs of SNV pairs involving consecutive nucleotides are highly correlated (r=0.98). Each point denotes an SNV-pair in one tumor sample. (**c**) VAFs of SNV pairs involving non-consecutive nucleotides exhibit a much lower correlation (r=0.27).

Other MSSGS result from the sequential acquisition of SNVs, as seen for *FGFR2*(S252W;Y375C), in which S252W appeared after neoadjuvant chemotherapy and both S252W and Y375C were found in a metastatic lymph node 2 years later, persisting in all nine subsequent samples (Fig. 4b, **Extended Data Table 1**). In yet other cases contributing SNVs made separate and sporadic appearances prior to detection of the complete MSSGS, as exemplified by an MSSGS involving *F8* in Subject 01 (Fig. 3a, **Extended Data Table 1**).

**Figure 4.**
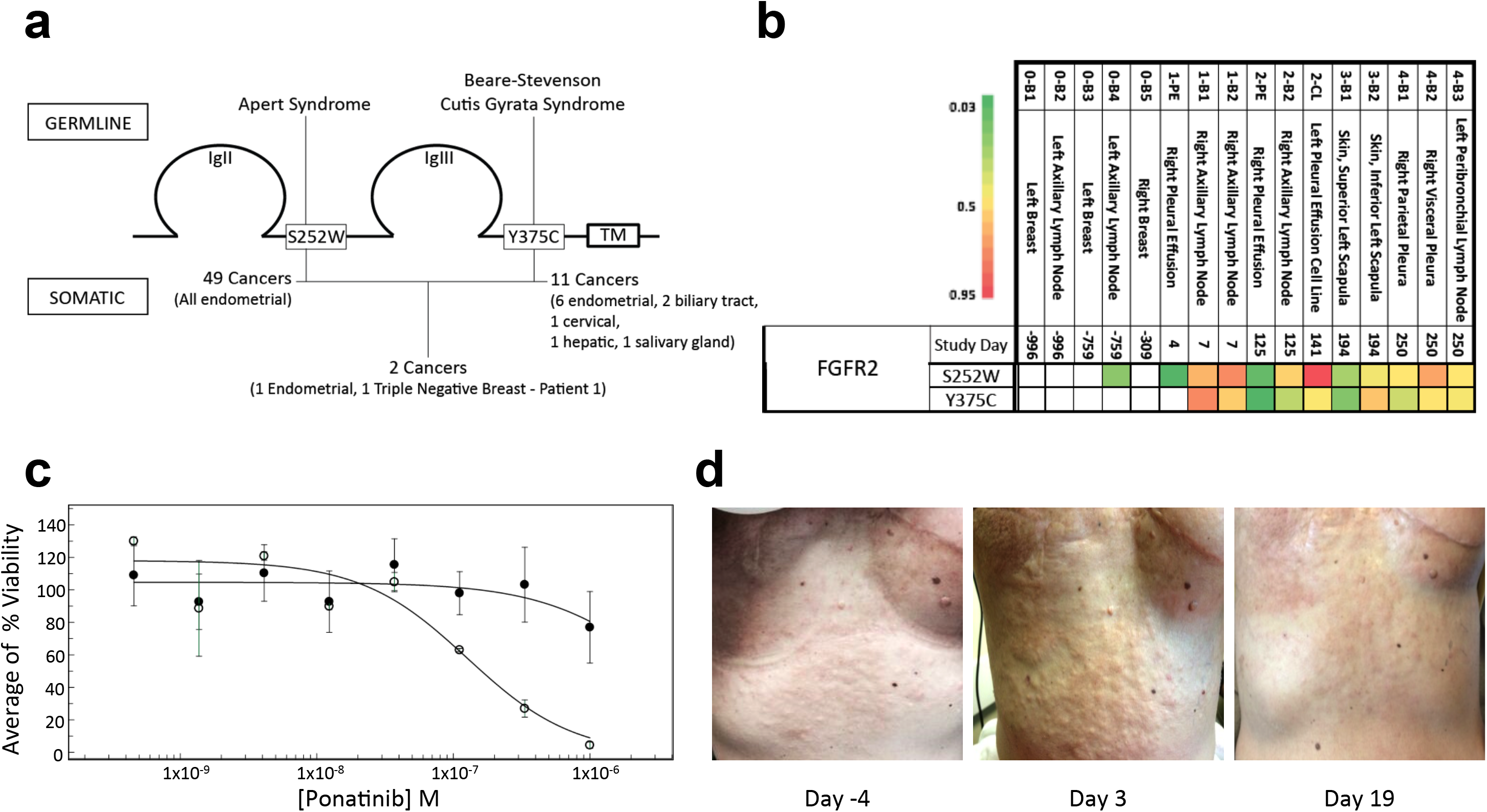
A cis-compound mutation drives TNBC progression. (**a**) Schematic depiction of two missense mutations affecting the FGFR2 extracellular domain. S252W and Y375C are well-characterized germline mutations that cause Alpert and Beare Stevenson Cutis Gyrata Syndrome, respectively. Both mutations have been identified previously in endometrial cancer (S252W - 49 cases; Y375C - 6 cases) whereas neither mutation has been reported previously in breast cancer. The same cis-compound mutation has been reported in a single case of endometrial cancer. (**b**) Appearance of point mutations encoding S252W and Y375C over the disease course of Subject 1. Note that S252W was acquired first, followed by Y375C. (**c**) Cell viability curves in response to a 3.5 log range of ponatinib concentrations. Closed circles show mean and standard deviations of 19 TNBC lines. Open circles indicate the response curve for transiently cultured breast cancer cells from Subject 1. (**d**) Clinical response to ponatinib in Subject 1, with resolution of breast cancer infiltrates in the skin.

Assessing allelic frequencies for all 426 MSSGS across 65 tumor samples revealed two distinct patterns. For the 71 MSSGS containing SNVs that affect consecutive nucleotides (including the above-referenced *ADAM33* and *ZZZ3*) we observed near perfect concordance in allelic frequencies between participating SNVs across tumor samples (Fig. 3b, **Extended Data Table 1**), consistent with their emergence from single mutational events (r=0.98). In contrast, concordance in allelic frequencies between SNVs separated by 1 or more nucleotides was significantly lower (r=0.27) (Fig. 3c, **Extended Data Table 1**). These findings indicate that the large majority of MSSGS containing immediately adjacent SNVs arose as single mutational events, whereas most other MSSGS were built from SNVs that arose separately.

Prior knowledge supports the functional significance of some MSSGS. For example, both missense mutations involving FGFR2(S252W;Y375C) (Fig. 4a) modify ligand binding^12^,^13^. FGFR2 S252W and Y375C have been separately associated with the congenital disorders Alpert Syndrome^14^ and Beare-Stevenson Syndrome^15^, respectively, and while each mutation has also been separately associated with endometrial cancer and FGFR2(S252W;Y375C) has been observed in a single case of endometrial cancer, neither mutation has been reported previously in breast cancer. To further evaluate *FGFR2*(S252W;Y375C) we extracted RNA from transiently cultured breast cancer cells from Subject 01, performed reverse transcriptase - PCR, and sequenced the cloned PCR fragments. Among 13 clones examined, 6 contained both mutations, 5 contained S252W only, and 2 were wildtype (data not shown). These results are consistent with the temporal acquisition of these SNVs, in which the S252W mutation arose first and Y375C was acquired later (Fig. 4b, **Extended Data Table 1**). A drug susceptibility screen using the same transiently cultured breast cancer cells demonstrated a much greater susceptibility to the FGFR2 inhibitor, ponatinib, compared to 19 TNBC cell lines (Fig. 4c), and ponatinib treatment in the patient produced a regression of breast cancer infiltrates in the skin (Fig. 4d). Possibly indicative of a growth advantage conferred by other MSSGS are results from Table 2, which show that for MSSGS involving *ABHD16Ac*, *TMPRSS13* and *ROBO4*, sequences bearing the entire cis-compound mutation are more abundant than sequences containing only one of the component SNVs.

We reasoned that if cis-compound mutations can confer a selective advantage in TNBC, MSSGS might be more prevalent than would be predicted from a random co-localization of SNVs. To address this question we evaluated 106 TNBC samples from the TCGA. Fig. 5s shows the number of SNVs contributing to MSSGS against the total SNV count for each of 106 TNBC samples from TCGA^4^. We also plotted a theoretical curve under the null hypothesis that MSSGS arise from the random co-localization of SNVs (see **Supplementary Methods**). The assumptions inherent in generating this theoretical curve exert a greater impact at high SNV counts; therefore, in Fig. 5b we exclude the 5 samples containing >600 SNVs, and focus on the remaining 101 samples carrying fewer than 200 SNVs. As expected, the number of SNVs contributing to MSSGS increases in accordance with the total number of SNVs. However, Fig. 5b also shows that the number of observed SNVs contributing to MSSGS tends to exceed levels predicted by the theoretical curve. These results indicate that for many samples, the number of SNVs contributing to MSSGS is higher than predicted had they arisen independently. To further examine this phenomenon, we performed a permutation test for each of the 106 patients to test the null hypothesis that SNVs co-localize to form MSSGS randomly (details are provided in Methods). With the 106 resulting p-values, we accept six patients when we use the Benjamini-Hochberg (BH) procedure^16^ to control the false discovery rate at 0.05 (Fig. 5b).

**Figure 5.**
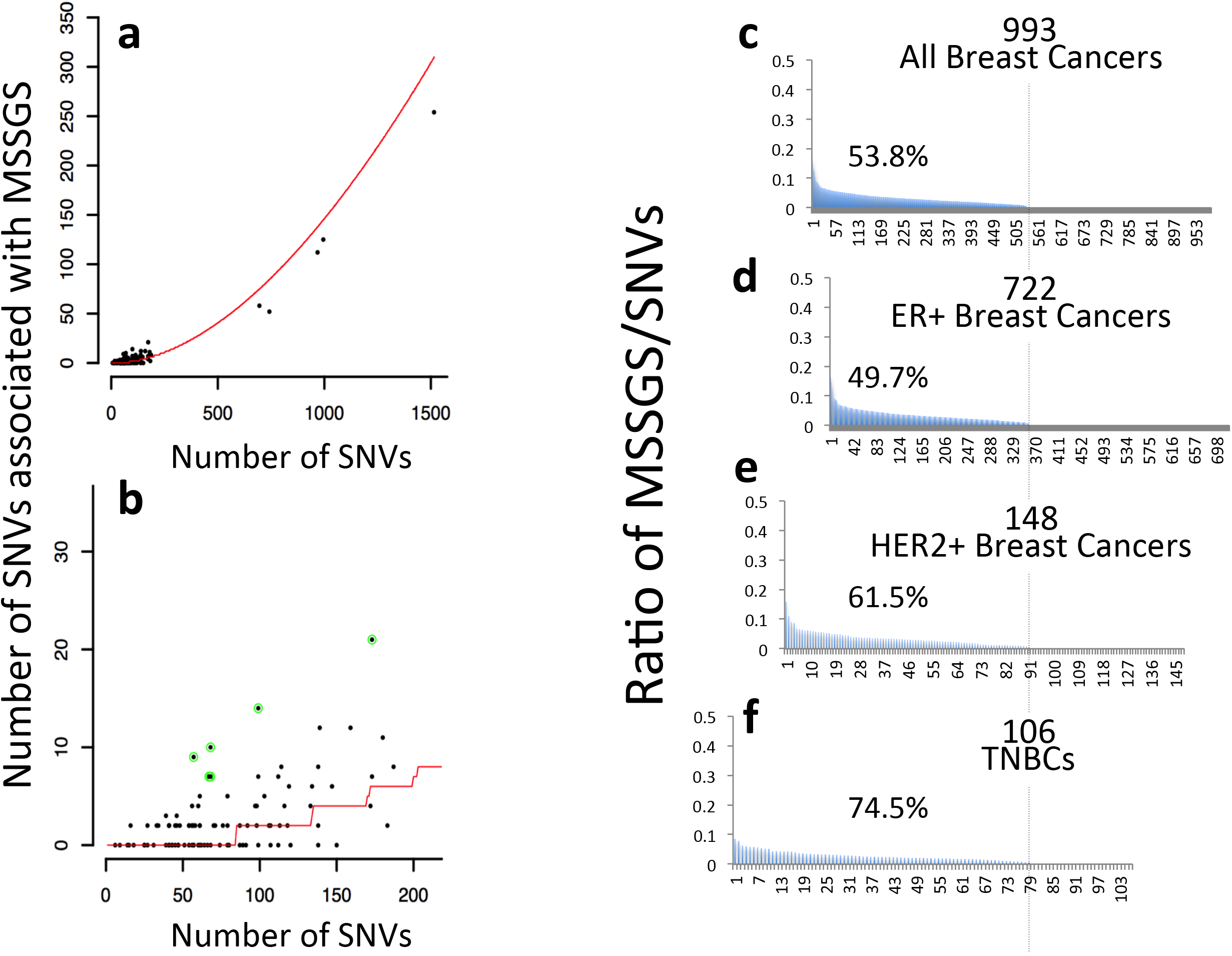
MSSGS arise at higher than predicted frequencies in TNBC. (**a**) Observed numbers of SNVs contributing to MSSGS (black dots) and theoretical numbers of SNVs contributing to MSSGS (red curve) against the observed numbers of SNVs in TCGA TNBC samples. **(b)** Same as **(a)** but magnified to show results for 101 TNBC patients with <200 SNVs. Green-circled dots are accepted by BH procedure at a FDR level of 0.05. **(c-f)** Ratios of MSSGS/SNVs for each of 993 breast cancer samples contained within TCGA, ordered from high to low. Vertical dashed line divides samples containing 1 or more MSSGS from samples that contain no MSSGS, and the percentages of samples containing at least one MSSGS is indicated. **(c)** all 993 breast cancer samples; **(d)** 722 ER+ breast cancer samples; **(e)** 148 HER2+ breast cancer samples; **(f)** 106 TNBC samples. Note that a higher proportion of TNBC samples contain at least one MSSGS.

Finally, we assessed the frequency of MSSGS in 993 breast cancer samples from TCGA. While there were no significant differences in the mean or median numbers of MSSGS per sample (data not shown), nor in the fraction of MSSGS per SNV, a higher percentage of TNBCs contained at least 1 MSSGS (74.5%) compared to HER2+ breast cancers (61.5%), or ER+ breast cancers (49.7%) (Fig. 5c-f).

Evidence presented here supports the involvement of MSSGS in tumor growth based on our findings that: i) The large majority of MSSGS are attributable to cis-compound mutations; ii) FGFR2(S252W;Y375C) has unambiguous functional significance; iii) sequences bearing the entire cis-compound mutation are frequently more abundant than sequences bearing only one of the component SNVs; and iv) MSSGS are detected at significantly higher frequencies than would be predicted from the random co-localization of SNVs in subsets of patients with TNBC. MSSGS lack many features of conventional driver mutations because they are not widely detected across tumors from different individuals, are typically present at sub-clonal levels, and are frequently not associated with cancer initiation but rather cancer progression. Some of the MSSGS described here are unlikely drivers of cancer progression, exemplified by *MUC4*, which resides in a region of increased mutation density (Extended Data Fig. 2).

Epistastic interactions between missense mutations can enable proteins to evolve new conformations and functions^17^,^18^. Cis-compound mutations associated with gain-of-function^19^,^20^ and loss-of-function^20^ underpin inherited disorders such as Multiple Endocrine Neoplasia Type 2B^19^, long QT syndrome^21^, and familial Alzheimer’s Disease^20^. Cis-compound mutations have also been described in cancer^22^^-^^32^, provide an important mechanism of resistance to BCR-ABL targeted tyrosine kinase inhibitors^29^^-^^31^, and can promote the re-acquisition of sensitivity in previously crizotinib resistant, ALK-rearranged lung cancer^32^. Driver mutations arising at metastatic sites of disease face a different set of constraints compared to driver mutations resident in primary tumors. Since cancers typically require years to become clinically evident, driver mutations conferring even a slight selective advantage can gain predominance due to the compounding effect of time^33^. In contrast, driver mutations acquired at metastatic sites of disease have a much shorter timeframe over which to gain a foothold, and the cells against which they compete have demonstrated a high degree of fitness. Therefore, driver mutations acquired at metastatic sites would be expected to be frequently present at subclonal levels. In conclusion, MSSGS appear to provide examples of “intramolecular epistasis” in the promotion of cancer progression^34^.

## METHODS

### Clinical Trial

ITOMIC-001 was approved by the Solid Tumor Scientific Review Committee and the Institutional Review Board of the Fred Hutchinson Cancer Research Center (FHCRC) and is registered at ClinicalTrials.gov (NCT01957514). Patients enroll from two sites: the Seattle Cancer Care Alliance (SCCA) and Northwest Medical Specialties (NWMS), a private practice with offices in Tacoma and Puyallup, Washington. Eligible subjects have metastatic TNBC, are platinum naive, and are scheduled to receive Cisplatin. Informed consent (Appendix 2) is conducted through detailed in person discussions involving, minimally, the principal investigator, the subject, and a subject family member, and last 1 - 2 hours. Study oversight is provided by a Data Safety Monitoring Board.

### Biopsies

Biopsies of metastases involving lymph nodes, subcutaneous tissues, and liver were performed under ultrasound guidance using an 18 gauge BioPince Full-Core Biopsy instrument. Up to 5 disease sites were biopsied in a single setting, and multiple biopsies were performed per disease site. Bone marrow biopsies were taken from the iliac crest using a Jamshidi T-Handle needle. Skin biopsies were performed using 3 - 4 mm punch biopsy instrument. In Subject 2 circulating tumor cells were collected by leukapheresis. All biopsies were performed using local anesthesia and most were performed using conscious sedation. Biopsy specimens were processed in accordance with a standardized set of operating procedures and subjects were contacted one day and one week following the procedure to assess for complications.

#### Rapid On-Site Evaluation

Core biopsy samples of metastases involving lymph nodes, soft tissues, or liver were immediately photographed and divided orthogonally. One half was formalin fixed (for formalin fixation and paraffin embedding [FFPE]) and the other half was gently pressed against an RNAse treated slide to generate a touch prep, then immediately snap frozen. Cytological evaluation of the touch prep was performed by a pathologist in real time and, if necessary, additional biopsies were procured to optimize sample size and tumor content. The time interval between sample acquisition and the completion of processing (known as the cold ischemia time) was recorded and was, except for the bone marrow biopsies and leukaphereses, less than 5 minutes in all cases (Table 2).

#### Selection of Biopsy Specimens for Further Testing

FFPE specimens were sectioned, stained with hematoxylin and eosin (H&E), and evaluated by a pathologist for tumor cell content and necrosis. FFPE samples judged to have sufficient tumor content from different metastatic sites were stained for HER2 and ER. To select samples for sequencing we examined H&E sections of the FFPE portion and touch preps of the corresponding snap frozen portion of each biopsy core, with the goal of maximizing tumor cell content. An uneven distribution of tumor cells across biopsy cores sometimes caused discordance in estimates of tumor cell content between the FFPE portion of a sample (assessed by morphology) and the corresponding snap frozen portion (estimated from the touch prep and variant allele frequencies from UW-OncoPlex).

#### Residual Clinical Materials

When feasible, leftover blood samples and pleural fluid specimens were processed and stored for analysis. The estimated time between sample collection and processing was recorded.

#### Autopsy Samples

At the time of this submission, Subjects 1, 2, 4, 5, 6, 7, 8, 11, and 12 had died and Subject 3 was lost to followup. Samples from Subjects 1, 2, 6, 11 and 12 were collected at the time of autopsy and processed as FFPE tissues."

#### DNA and RNA extraction

DNA and RNA were extracted from samples using AllPrep DNA/RNA/Protein Mini Kit from Qiagen following the manufacturer’s protocol. The quality of the RNA was analyzed using the Agilent 2100 Bioanalyzer System.

### Whole Exome Sequencing

WES is performed by the UW Northwest Clinical Genomics Laboratory, a CLIA--certified facility at the UW. DNA fragment libraries were constructed from tumor biopsies and blood (normal) using the Hyper Prep Library Preparation Kit (KAPA Biosystems, Wilmington, MA). Each library was enriched for protein and RNA coding portions of the human genome using the SeqCap EZ Exome v3.0 (Roche NimbleGen, Madison, WI) capture system. The target includes all coding content from the CCDS, RefSeq and miRBase databases. Paired-end sequencing (100 bp) of enriched libraries was performed using TruSeq Rapid Sequencing-by-Synthesis chemistry on a HiSeq 2500 (Illumina, San Diego, CA) sequencer according to the manufacturer’s recommended protocol. Base calls were generated in real-time on the HiSeq instrument (RTA 1.17.21.3 software). After sequencing was complete, resulting reads were demultiplexed and sample-specific FASTQs produced. The average depth of coverage was 188 and the range was 166 to 208. The resulting reads are aligned to the genome human reference (hg19) using BWA (Burrows-Wheeler Aligner), duplicate reads removed with Picard and variants called with GATK (Genome Analysis Toolkit). BAM files are uploaded to DNAnexus (Mountainview, California) for subsequent somatic variant detection by researchers at UC Santa Cruz, UW and Data4Cure (using a proprietary algorithm).

### Whole Genome Sequencing

Confirmation of a subset of MSSGs was done by NantOmics (Culver City, CA), which performed whole genome sequencing (WGS) in 39 of the 67 samples described here, using the same DNA that had been tested by WES. WGS Sequencing was performed on the Illumina HiSeq X sequencing platform using libraries prepared via the KAPA Hyper prep kit. Tumor genomes were sequenced to an average depth of 60x, and Normal genomes were sequenced to an average depth of 30x. Mutations were identified using the CLIA-validated NantOmics Contraster pipeline as previously described^35^.

### Germline Genome

Germline variant detection is performed to identify genes associated with inherited cancer syndromes, such as mutations involving *BRCA1* or *BRCA2*.

### Data Management

Clinical data are entered into REDCap, a web--based application for Electronic Data Capture. The Institute of Translational Health Science’s installation of REDCap is hosted on a secure server at UWMC. Large datasets that are not well suited for storage on REDCap are stored on the DNANexus platform (https://platform.dnanexus.com).

### Clonal Analysis of MSSGS

MSSGS-containing DNA fragments were PCR-amplified using primers flanking all participating SNVs. The amplified PCR fragments were cloned into pBluescript II KS+ vector and transformed into *E. coli*. For each MSSGS, DNA from about 100 colonies was individually prepared and sequenced, and the number of clones containing germline sequence or one or multiple participating SNVs was determined.

### Comparing sizes of genes affected by MSSGS versus isolated SNVs

To assess whether the formation of MSSGS is related to gene size, we compared the distributions of the median transcript sizes between genes affected by MSSGs versus genes affected by isolated SNVs. We performed the Wilcoxon rank-sum test to compare the means of the two distributions and found no significant difference (Extended Data Fig. 1). Gene transcript sizes were obtained from Biomart (Ensembl).

### Haplotype phasing of SNVs associated with MSSGS

To assess the orientation of neighbouring SNVs within MSSGs, we used ReadBackedPhasing from Genome Analysis Toolkit version 3.3.0. Phasing information with at least 20.0 quality score was used to assess whether a pair of SNVs are cis or trans.

### Evaluating the local mutation rates in regions surrounding MSSGS vs isolated SNVs

To assess whether the formation of MSSGS is driven by elevated local mutation rates, we compared local mutation rates surrounding MSSGS with mutation rates genome wide. In order to calculate the local mutation rate for each MSSGS, we counted the number of SNVs identified by WGS in a 1 megabase window centered on each MSSGS (obtained by subtracting transcriptional stop site by the transcriptional start site, using RefSeq transcripts as a reference). We then calculated the local mutational rate across the genome by windowing each sample into 1MB windows and counting the number of SNVs. A plot of the local mutation rates surrounding MSSGS is depicted by the red circles shown in Figure 2a. Additionally, the distance between each neighboring SNV identified in the WGS samples was calculated to identify regions of kataegis^9^, which we defined as at least 10 SNVs in a 100 kilobase window. Only one MSSGS, involving *MUC4*, landed in such a region (Figure 2b).

### The theoretical distribution of SNVs contributing to MSSGS

Suppose that a patient carries m SNVs in the tumor cells. Let N (N=17,672) be the total number of genes subject to possible mutation in TNBC, and W be a vector of length N. We assume that each gene can be mutated with a probability proportional to W, and W = 0.8W_1_ + 0.2W2, where W_1_ is the normalized gene mutation frequency in TNBC samples in TCGA and W_2_ is the normalized gene mutation frequency in non-TNBC BC samples in TCGA. We include the term 0.2W_2_ because there are only 106 TNBC samples (in which 7,933 genes are mutated), and we want to use W_2_ to mimic the unseen genes (which can be potentially mutated in TNBC). In each round of permutation, we randomly assign each of the *m* SNVs to one of the 17,672 genes with a probability proportional to W, and then count SNVs in MSSGS. We repeat this procedure 10,000 times, and each time we obtain a count of the number of SNVs contributing to MSSGS. We then use the median of these 10,000 counts as the expected number of SNVs contributing to MSSGS. The entire simulation procedure is repeated for m from 1 up to 1500, where 1500 is the maximal number of SNVs observed in the TCGA cohort.

To assign a patient-specific P-value, we repeat the above simulation procedure using the observed value of m. We then compare the observed count of SNVs contributing to MSSGS with the 10,000 count values from the simulation. We define the P-value as (t + 1)/10001, where t is the number of times that the observed count of SNVs contributing to MSSGS is less than or equal to the count from simulations.

### High Throughput Drug Screen

The drug sensitivity profiles of transiently cultured cells from a pleural effusion from Subject 1 and 19 TNBC lines were compared in a high throughput screen of 160 approved and investigational oncology drugs. Cells were seeded into non-tissue culture-treated 384-well plates. Twelve hours later, compounds were added and after 72 hours viability was assessed. Resulting dose curves were fitted using idbs’ XLFit and to a 4 Parameter Logistic Dose Response Model. The TNBC lines tested were BT-20, BT549, HCC1143, HCC1187, HCC1395, HCC1599, HCC1806, HCC1937, HCC2157, HCC38, HCC70, Hs 578T, MDA-MB-157, MDA-MB-436, MDA-MB-231, MDA-MB-453, MDA-MB-468, MFM-223 and SW-527.

## ACKNOWLEDGEMENTS

The patients and their families. We also gratefully acknowledge the assistance of Michelle Flores, Melodee Smith, Sherry Littlefield, Laura Connelly-Smith, James Riddle, Ednus Warren, Stan Riddell, Julie Gralow, Bernie McLaughlin, Jennifer Specht, Brent Wood, Katie Dougherty, Anju Thomas, Paul Martin, Corinne Fligner, David Yadock, Jennifer Spokely, Krystal Malhotra, Stephanie Parker, Daniel Campton, Josh Nordberg, Nick Drovetto, Nicole Heying, Leila Ritter and NW BioTrust. This publication was supported by the National Center for Advancing Translational Sciences of the National Institutes of Health under Award Number UL1TR000423. Funding for this work was provided by South Sound CARE, the Gary E. Milgard Family Foundation, the Pano Koumantaros Cancer Research Fund, the Washington Research Foundation, the University of Washington School of Medicine, and private donors.

## AUTHOR CONTRIBUTIONS

N.H., J.L., S.R.S., A.J.R., M.G., J.M.J., V.K., M.L-M, S.C.B., D.H., J.Z., W.L.R., W.S.N. and C.A.B. contributed to data analysis and interpretation. C.Z., J.G., M.O.D., C.C.P., S.C.S., S.R.S., A.J.R., S.H., and C.A.B. contributed experimental results, K.A.B., S.B., F.M.S., W.L.M., S.P., S.K.A., V.K.G., and P.S.-S. contributed to the clinical trial, and N.H., J.L., Z.D., W.L.R., W.S.N. and C.A.B. wrote the manuscript.

## LEGENDS FOR EXTENDED DATA FIGURES AND TABLES

**Extended Data Figure 1.** CDF plots depicting distributions of median transcript sizes for genes affected by MSSGs versus genes affected by isolated SNVs. Results indicate that there is no significant relationship between gene size and MSSG formation.

**Extended Data Table 1.** Summary of genes bearing multiple SNVs across 67 tumor samples. Rows indicate gene names, nucleotides at reference and alternate alleles, consequence, position and amino acid substitutions of missense mutations, codon changes, predicted functional consequences using Sift and Polyphen, and chromosomal positions. Columns indicate sample description, number of days post diagnosis, study day, whether tissue was snap frozen or stored as formalin fixed paraffin embedded tissue (FFPE) or both, cold ischemia time, and estimated tumor purity. 3 columns describe each sample (designated 01-0-B1, etc). Ref = number of reads for reference allele; Alt = number of reads for alternate allele; VAF = variant allele frequency. Blank cells denote a value of zero.

**Extended Data Table 2. Confirmation of MSSGS by WGS**. To confirm MSSGS identified by WES, 39 samples evaluated by WES were also examined using WGS. Results for each sample and MSSGS-associated SNV are shown. WGS_Ref and WGS_Alt indicate read counts for MSSGS-associated SNVs identified independently by WGS. WGS_RAW_Depth and WGS_RAW_ALT indicate read depths for MSSGS-associated SNVs identified by WES but not independently identified by WGS. P values were calculated using Fisher’s exact test.

**Extended Data Table 3.** Haplotype phasing^10^ reveals that 394 of 407 MSSGS accessible for analysis (97%) co-localized to the same allele.

## ACKNOWLEDGEMENTS

We acknowledge foremost the patients and their families. We also gratefully acknowledge the assistance of Michelle Flores, Melodee Smith, Sherry Littlefield, Laura Connelly-Smith, James Riddle, Ednus Warren, Stan Riddell, Julie Gralow, Bernie McLaughlin, Jennifer Specht, Paul Martin, Brent Wood, Katy Dougherty, Anju Thomas, Corinne Fligner, David Yadock, Jennifer Spokely, Krystal Malhotra, Zhijun Duan, and NW BioTrust. We thank Kate Sweeney for assistance with figure preparation. This publication was supported by the National Center for Advancing Translational Sciences of the National Institutes of Health under Award Number UL1TR000423.

